# BBBP-Atlas: Unified Interpretable Modeling of Blood–Brain Barrier Permeability across Small Molecules and Peptides

**DOI:** 10.64898/2026.07.06.736742

**Authors:** Xin Shen, Qun Su, Hao Luo, Qiaolin Gou, Jingxuan Ge, Tingjun Hou, Jike Wang, Yu Kang

## Abstract

Accurate prediction of blood-brain barrier permeability (BBBP) is essential for central nervous system drug discovery, yet existing models are often limited by their reliance on predefined physicochemical descriptors, small-molecule-centered training sets, or conformation-dependent representations, which restricts their transferability across chemically diverse modalities especially peptides. In addition, publicly available BBBP datasets remain fragmented, inconsistently standardized, and weakly controlled for molecular redundancy, increasing the risk of data leakage and overestimated model performance. In this study, we propose BBBP-Atlas, a structure-aware BBB permeability prediction model designed for unified modeling of small molecules and peptides with the first cross-modal dataset OmniBBBP. Designed to bypass descriptor and conformation dependencies, our model represents standardized molecular structures as atom-level graphs to capture local atom-bond environments and long-range topological dependencies associated with BBB transport. This design enables direct learning of structure-permeability relationships from molecular topology. For model training and evaluation, we curated a cross-modal, redundancy-filtered database OmniBBBP that seamlessly unifies small molecules and complex peptides, containing 10,218 unique compounds with 9,316 small molecules and 902 peptides. BBBP-Atlas achieved an accuracy of 0.8914 and an MCC of 0.7678 on the independent test set. On a balanced external benchmark of 200 compounds, our model reached an AUC of 0.9108, an accuracy of 0.8500, and an MCC of 0.7000, outperforming LightBBB by an absolute MCC gain of 6%. Case studies further showed that BBBP-Atlas captured clinically meaningful BBB permeability patterns, correctly identifying lorlatinib as BBB-permeable and vancomycin as BBB-impermeable with high confidence. The OmniBBBP-backed BBBP-Atlas offers a versatile and cross-modal approach for single-compound prediction, batch screening, and dataset exploration for CNS drug discovery. BBBP-Atlas is available at https://cadd.drugflow.com/bbbp/.

## INTRODUCTION

Central nervous system (CNS) drug discovery remains particularly challenging due to the presence of the Blood-Brain Barrier (BBB), a specialized vascular interface that tightly regulates molecular exchange between the bloodstream and the brain.^1^ The BBB is primarily formed by brain microvascular endothelial cells connected by tight junctions and supported by pericytes and astrocytic end-feet, collectively forming a highly selective neurovascular unit that restricts molecular entry into the brain.^2,3^

Although this barrier is essential for maintaining neural homeostasis, it also severely limits the delivery of therapeutic agents. Nearly 98% of small-molecule drugs and almost all large-molecule therapeutics fail to penetrate the BBB effectively.^4^ Consequently, CNS drug development has one of the lowest success rates among therapeutic areas, with only approximately 6.2% of candidates advancing from clinical trials to regulatory approval.^5^ Poor brain exposure resulting from inadequate BBB penetration is widely recognized as a major cause of CNS drug failure, highlighting the critical need for early and reliable assessment of BBB permeability.

BBB permeability is governed by a delicate balance between intrinsic molecular physicochemical properties and biological transport mechanisms. Passive diffusion is primarily governed by molecular weight (MW),^6^ lipophilicity (LogP),^7^ and hydrogen-bonding capacity,^8^ while active transport systems such as efflux pumps^9^ further modulate brain exposure. Early predictive efforts focused on quantitative structure-activity relationship (QSAR) models using empirical rules and physicochemical descriptors, including polar surface area (PSA), molecular weight, and lipophilicity. For example, Young et al. reported a strong correlation between logBB and Δ logP, defined as logP(o/w)-logP(cycl/w),^10^ while Kaliszan and Markuszewski provided a physicochemical interpretation linking increased BBB permeability to higher lipophilicity and smaller molecular size.^11^ Despite these insights, traditional QSAR models are limited by descriptor redundancy and susceptibility to overfitting, which restricts their generalizability across diverse chemical scaffolds.^12^

To overcome these challenges, subsequent studies integrated richer feature representations and applied machine-learning algorithms, including support vector machines (SVM)^13^ and gradient boosting^14^, to enhance model robustness and generalization. Lee et al. explored explainable machine-learning frameworks that capture synergistic effects among molecular substructures in BBB permeability prediction.^15^ More recently, deep learning approaches have been introduced to directly learn molecular representations from graph structures.^16–18^ For instance, Nguyen et al. proposed the geometric multi-color message-passing neural network (GMC-MPNN),^19^ which incorporates atomic-level geometric features and long-range interactions to improve generalization. Although GMC-MPNN achieves the AUC-ROC scores ranging from 0.921 to 0.947 on standard small-molecule benchmarks, its applicability remains inherently tied to the availability of high-quality 3D structural conformations. In addition, it has yet to be generalized to the complex peptide systems featuring non-natural or chemically modified residues.

Beyond model development, expanding dataset scale has also been explored to improve chemical space coverage. For instance, Shaker et al. compiled a dataset of 7,162 compounds^20^ to support large-scale BBB permeability modeling. However, simply increasing data quantity without rigorous curation often introduces noise rather than informative signal, leading to models that generalize poorly to novel chemical scaffolds. Despite these advances, several fundamental limitations persist. First, many publicly available datasets contain substantial redundancy (up to 30%), where different non-standardized Simplified Molecular Input Line Entry System (SMILES) correspond to identical molecules,^20^ leading to data leakage and inflated performance estimates. Second, most existing models rely primarily on two-dimensional physicochemical descriptors, which fail to capture structural features relevant to BBB transport. Third, many studies focus exclusively on either small molecules or peptides, limiting applicability across diverse molecular modalities. Consequently, models may achieve high accuracy by exploiting dataset-specific correlations rather than learning transferable structure–permeability relationships. Together, these limitations raise a fundamental question: to what extent can BBB permeability be reliably inferred from intrinsic molecular structure alone, particularly across chemically distinct modalities such as small molecules and peptides?

To address these limitations, we present BBBP-Atlas, a unified structure-aware framework powered by a customized topology-driven architecture, alongside OmniBBBP, the world’s first cross-modal, leakage-free reference benchmark for BBBP prediction. Developed to entirely bypass traditional descriptor and conformation dependencies, the proposed model translates standardized molecular structures into atom-level graphs. This architecture is specifically engineered to capture both local atom-bond environments and long-range topological dependencies associated with BBB transport, thereby enabling the direct extraction of structure-permeability relationships solely from molecular topology. To power the training and evaluation of our proposed model, we established the largest and the first cross-modal BBBP database to date. OmniBBBP contains 10,218 unique compounds (9,316 small molecules and 902 peptides) collected from 74 public sources, making it five times larger than the widely used MoleculeNet^21^ BBBP dataset and the largest publicly available collection of peptide BBB data.^22^ Comprehensive benchmarking evaluations demonstrate that our tailored architecture consistently outperforms state-of-the-art (SOTA) baselines, yielding substantial performance gains in prediction accuracy and Matthews correlation coefficients (MCC) across both independent and external balanced tests. All data and the pretrained model are freely accessible through an interactive web platform (https://cadd.drugflow.com/bbbp/). Beyond single-compound and high-throughput batch prediction, this open-access platform uniquely supports intuitive dataset exploration. By successfully bridging the gap between small molecules and complex peptide modalities, our BBBP-Atlas framework provides a highly standardized, non-redundant, and structure-aware benchmark pipeline, offering a robust and extensible foundational tool for computational CNS drug discovery.

## RESULTS

### Overview of BBBP-Atlas

Aiming for a unified understanding across molecular modalities, we constructed OmniBBBP, a refined dataset integrating experimentally validated permeability annotations for both small molecules and peptides. After molecular standardization and preprocessing using RDKit (**Figure 1a**), the dataset comprises 10,218 unique compounds, including 9,316 small molecules and 902 peptides (**Supplementary Table 1**) collected from 74 sources. Among these, 1,574 entries include experimentally measured logBB values, providing quantitative estimates of brain exposure. This dataset expands the coverage of peptide-related BBB data, which has been underrepresented in previous studies. Despite its broad structural diversity, dimensionality reduction analyses (LDA and PCA) reveal a substantial overlap in chemical space between permeable and non-permeable compounds, indicating that linear decision boundaries are insufficient to separate the two classes (**Supplementary Figures 2-3**).

**Figure 1.**
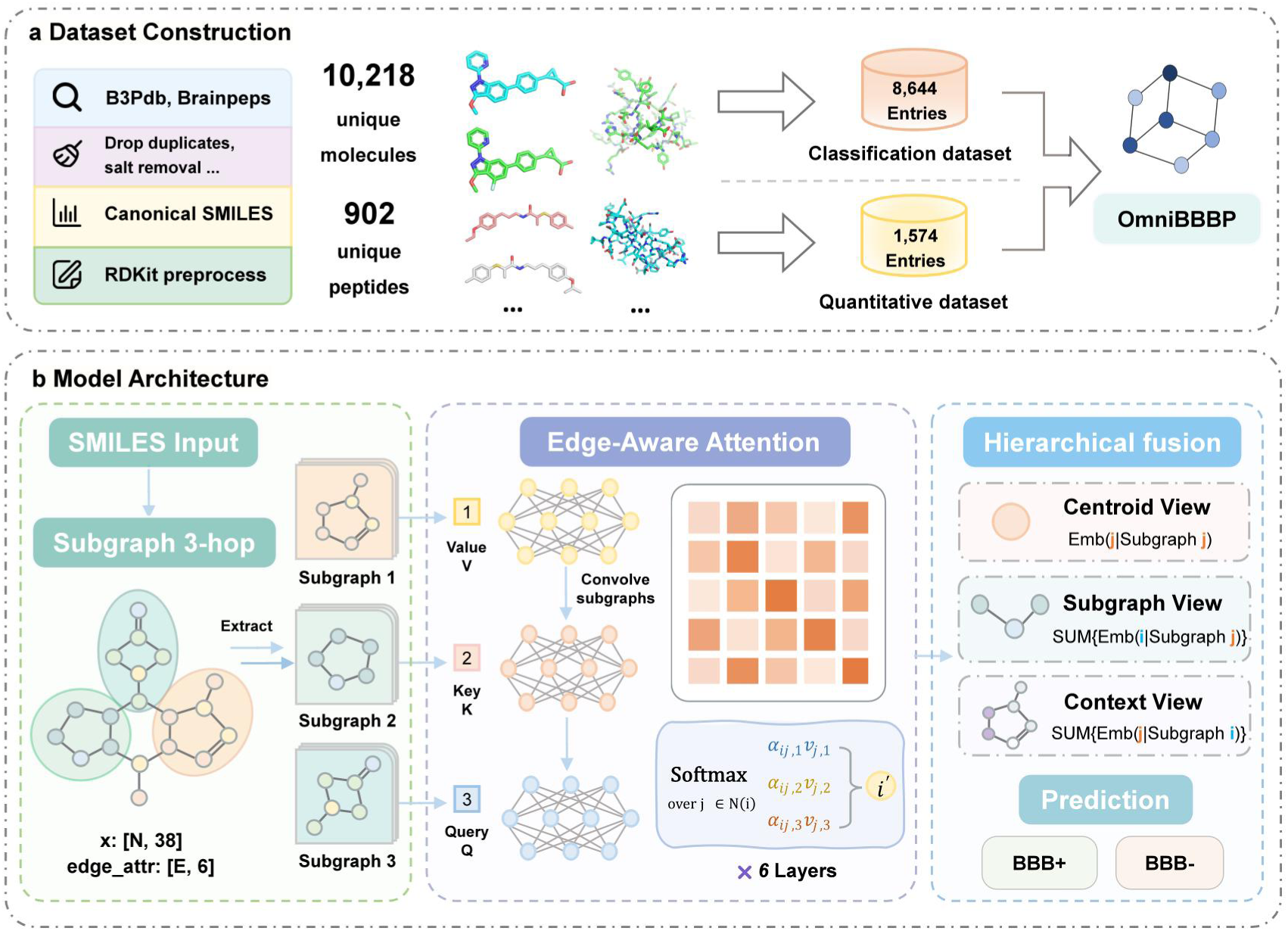
Overview of the BBBP-Atlas framework. (a) OmniBBBP dataset curation. Multi-source integration across 74 raw repositories to assemble the world’s first cross-modal, leakage-free benchmark containing 10,218 unique therapeutic entities via RDKit-based standardization and strict molecular redundancy filtering. (b) BBBP-Atlas model. Descriptor-free embedding layer 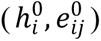 converting molecular chemistry into atom-level graphs, coupled with stacked message-passing and tailored multi-head self-attention mechanisms to dynamically capture multi-scale, long-range topological dependencies across diverse chemical modalities.

Considering the structural complexity and heterogeneity of the BBBP chemical space, we proposed a novel architecture BBBP-Atlas (**Figure 1b**) designed to capture long-range atomic dependencies and higher-order structural features. Rather than relying on fixed global descriptors, our model treats each compound as an attributed graph to evaluate distal substituents, ring systems, and peptide backbone motifs in a context-dependent manner via a self-attention mechanism. As messages propagate through stacked transformer layers, localized chemical environments are progressively integrated into molecule-level embeddings. This representation places compact small-molecule scaffolds and extended peptide-like structures into a shared latent space, bridging the rigid scaffolds of small molecules and the flexible, repeating backbones of peptides.

Furthermore, we incorporated model interpretability to align model decisions with established chemical intuition through SHAP and counterfactual analysis. To support practical application, the dataset and pretrained models are deployed through a web platform (https://cadd.drugflow.com/bbbp/). The platform supports single-compound prediction from SMILES input and batch processing via CSV upload (**Supplementary Figure 4**). Integrated database and statistics modules further empower users to explore chemical space and physicochemical property distributions, serving as a transparent and robust utility for virtual screening and candidate prioritization.

### Model performance benchmarking

To evaluate the predictive capability of the proposed framework, we conducted benchmark comparisons on the BBBP-Atlas dataset. The representative baselines include AttentiveFP,^23^ XP-GCN,^24^ BBBP_MPNN, DeepBBBP,^16^ as well as classical machine learning models (ExtraTrees, Gradient Boosting, and Bayesian-optimized XGBoost).

Given that many public BBB permeability datasets are confounded by inconsistent molecular representations and redundant entries arising from alternative SMILES encodings, which inevitably induce data leakage and performance inflation. To ensure fair and reproducible comparisons, all models were trained and evaluated on the standardized, deduplicated BBBP-Atlas dataset using an 8:1:1 train/validation/test split. Performance metrics (**Supplementary Table 2**) included accuracy (ACC), F1 score (F1), Matthews correlation coefficient (MCC), sensitivity (SE), specificity (SP), and balanced accuracy (BA), all computed on the held-out test set.

As summarized in **Figure 2**, our model achieves an MCC of 0.7678, comparable to DeepBBBP (0.7595) and XP-GCN (0.7451), indicating a competitive and unbiased benchmark. Our curation process removed redundancies, reducing the risk of inflated performance estimates.

**Figure 2.**
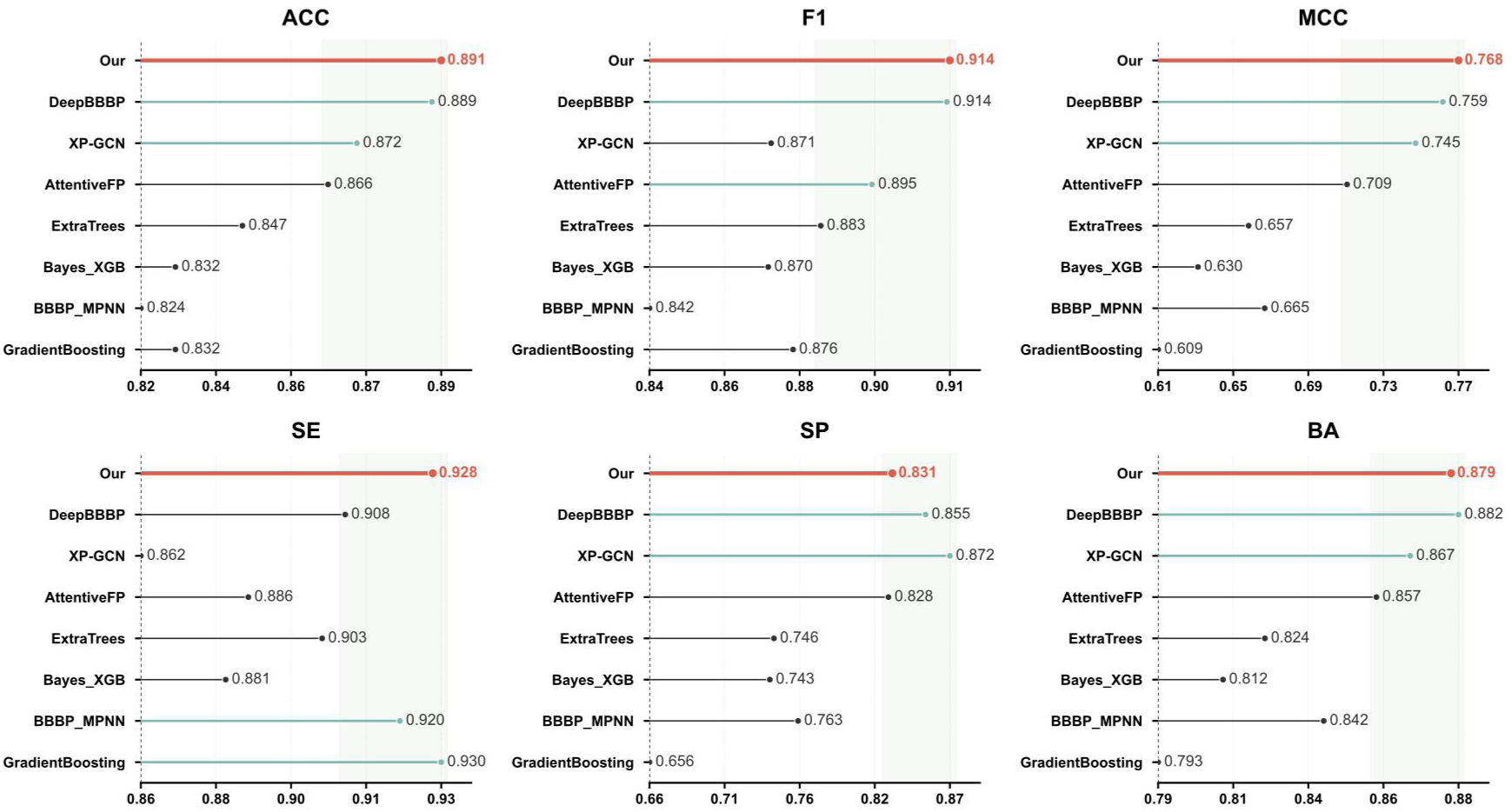
Benchmark performance on the independent test set (8:1:1 split). Performance comparison of eight models evaluated on the independent test set using ACC, F1, MCC, SE, SP, and BA. The proposed model consistently yields superior or competitive results across most metrics, indicating strong generalization capability under standard data splitting.

Our geometry-informed network balances sensitivity and specificity, a critical consideration in CNS drug discovery. In early-stage discovery pipelines, false negatives are exceptionally detrimental as they prematurely eliminate potentially viable neuroactive scaffolds, whereas false positives can be filtered through downstream experimental validation. The model manifests high sensitivity (SE = 0.9279) while maintaining competitive specificity (SP = 0.8309). This balance identifies permeable compounds effectively while limiting false positives. In comparison, Gradient Boosting achieves similar sensitivity (0.9298) but exhibits substantially lower specificity (0.6557), leading to an elevated false-positive rate. Conversely, DeepBBBP prioritizes specificity (0.8549) at the expense of sensitivity (0.9082), potentially missing viable candidates.

BBBP-Atlas provides the most mathematically robust and biologically practical framework for early-stage high-throughput CNS drug screening.

### Benchmark and out-of-distribution generalization

To further assess out-of-distribution (OOD) generalization, we used a scaffold-based split to build an external test set structurally dissimilar from the training set, mimicking real-world prospective screening. The set was balanced with 100 BBB-permeable (BBB+) and 100 non-permeable (BBB-) compounds (**Supplementary Table 4**) to eliminate class-imbalance bias.

We prioritize the MCC for cross-model comparison, as it penalizes skewed sensitivity-specificity trade-offs that ACC or F1 might obscure^25,26^. Even on a balanced dataset, ACC and F1 can mask differences in error distribution across confusion matrix components. Models with similar ACC and F1 may exhibit markedly different sensitivity-specificity trade-offs, which MCC captures more reliably. On the small-molecule set (**Table 1**), our model achieved a MCC of 0.700, outperforming all baselines (**Figure 3**). While BBBP_MPNN achieved a comparable ACC (0.815), it severely sacrificed sensitivity (SE = 0.71) for high specificity (SP = 0.92) (**Supplementary Figure 4**). In practical CNS drug discovery, such a bias yields a 30% false-negative rate, prematurely rejecting viable therapeutic candidates. In contrast, our model maintained well-balanced predictive performance (SE = 0.85, SP = 0.85), underscoring its reliability for unbiased screening.

**Figure 3.**
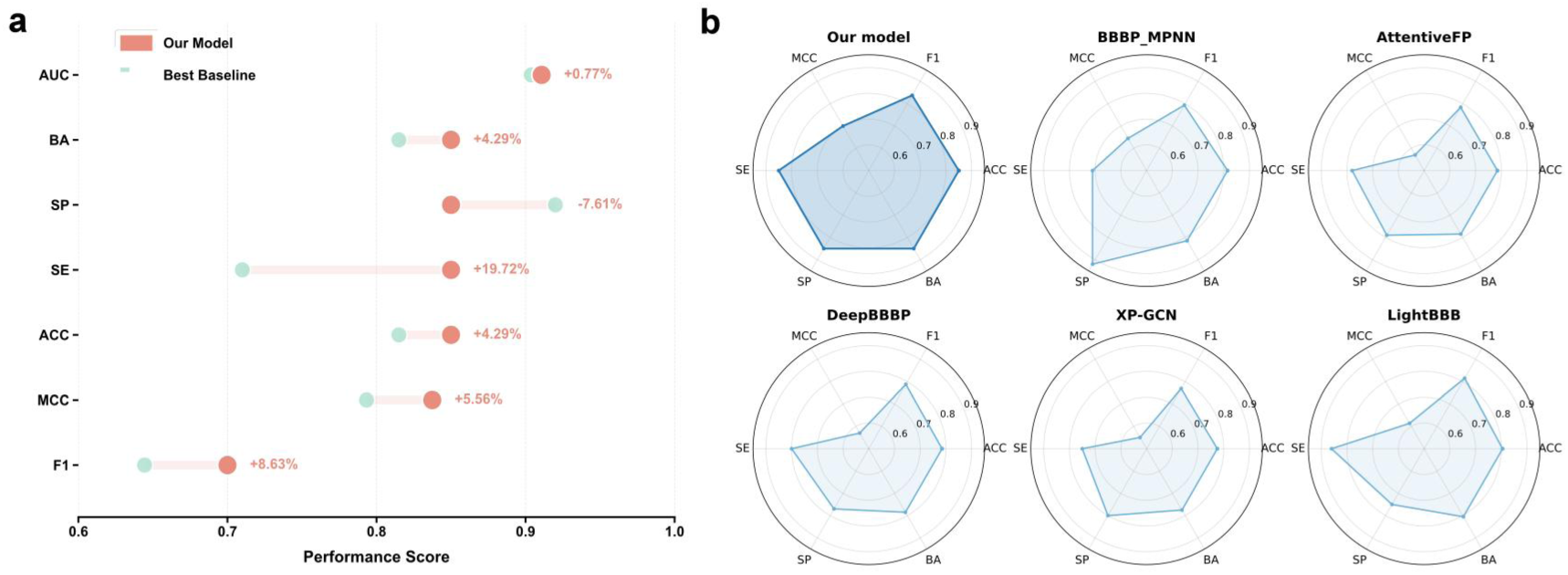
Benchmark performance on the external benchmark dataset. (a) Model gains compared to the best-performing baseline using AUC, BA, SP, SE, ACC, and MCC. Percentage values indicate the relative performance improvement or decline of the proposed model compared with the corresponding baseline. (b) Radar-chart comparison of representative models using ACC, F1, MCC, SE, SP, and BA. The proposed model exhibits a more balanced metric profile and improved robustness on unseen external data, supporting its generalization capability beyond the training distribution.

**Table 1.** Performance Comparison on an Independent, Balanced External Test Set of Small Molecules. All models were evaluated on the same independently sampled external test set with balanced class distribution. Performance metrics (AUC, MCC, ACC, F1-score, SE, SP, BA) were calculated to ensure a fair comparison.

| Model | AUC | MCC | ACC | F1 | BA | SE | SP |
| --- | --- | --- | --- | --- | --- | --- | --- |
| Our model | 0.9108 | 0.7000 | 0.8500 | 0.8374 | 0.8500 | 0.8500 | 0.8500 |
| BBBP_MPNN | 0.9038 | 0.6444 | 0.8150 | 0.7933 | 0.8150 | 0.7100 | 0.9200 |
| AttentiveFP | 0.8862 | 0.5700 | 0.7850 | 0.7839 | 0.7850 | 0.7800 | 0.7900 |
| DeepBBBP | 0.8906 | 0.5703 | 0.7850 | 0.7882 | 0.7850 | 0.8000 | 0.7700 |
| XP-GCN | 0.8641 | 0.5507 | 0.7750 | 0.7692 | 0.7750 | 0.7500 | 0.8000 |
| LightBBB<sup>20</sup> | / | 0.6137 | 0.8050 | 0.8152 | 0.8050 | 0.8600 | 0.7500 |

A zero-shot transfer of models trained solely on small molecules to peptides yielded near-random performance (ACC ≈ 0.58), indicating a large domain shift. From a biophysical perspective, peptides possess larger solvent-accessible surface areas, greater conformational flexibility, and more intricate hydrogen-bonding networks, features that likely raise the energy barriers for BBB penetration in a non-linear manner.

On a mixed OOD dataset, however, the unified model retained stable performance (ACC = 0.779, MCC = 0.564) as shown in **Figure 4**. This illustrates that the cross-modal framework captures partially shared pharmacophoric determinants, such as topological polar surface area and amphiphilicity, mapping structurally diverse compounds into a common latent space. However, the persistent performance gap between modalities (**Supplementary Table 3**) highlights the limitations of a one-size-fits-all approach under substantial domain shift, indicating that future improvements may benefit from incorporating modality-specific structural priors.

**Figure 4.**
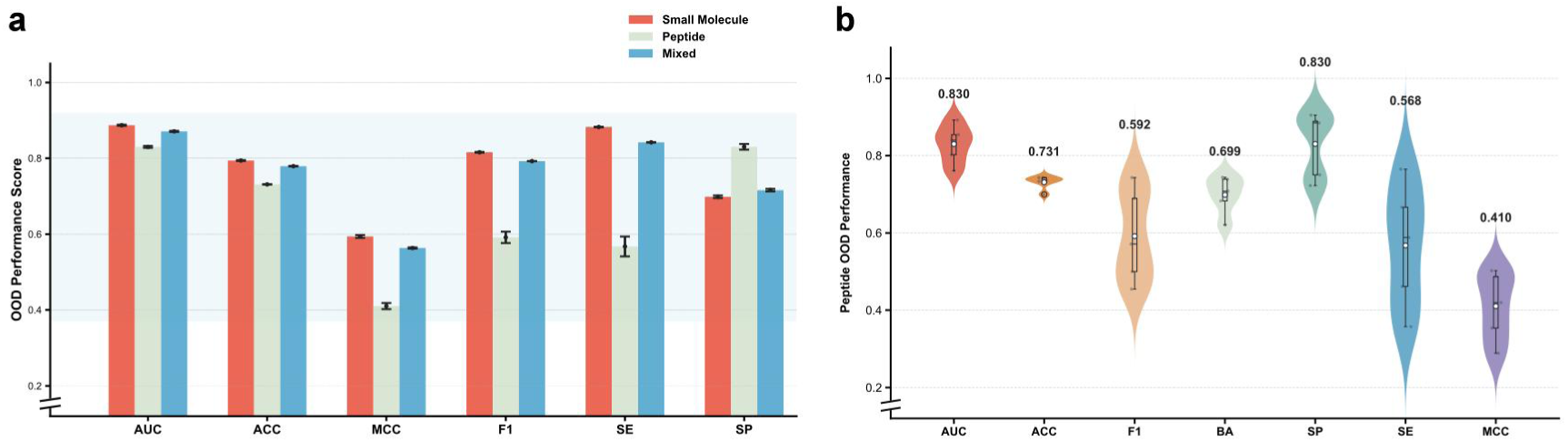
Cross-modality performance across out-of-distribution (OOD) scenarios. (a) Performance on three external test sets (Small Molecule, Peptidomimetic, and Mixed) reveal the modality gap and a marked decline on peptides especially in MCC and SE. However, the proposed model maintains robust cross-modality performance and competitive generalization relative to traditional single-modal approaches. (b) Evaluation on peptide-only datasets demonstrates the model’s capacity to capture transferable representations across distinct chemical spaces, resulting in enhanced predictive accuracy and generalization.

### Model interpretability analysis

To elucidate the decision-making process of BBBP-Atlas, we employed SHapley Additive exPlanations (SHAP)^27^ for global feature attribution (**Figure 5**), with local counterfactual (CF) analysis. These CF pairs characterize the minimal structural perturbations required to invert predictions,^28,29^ providing a localized view of how specific modifications influence model decisions across diverse molecular contexts.

**Figure 5.**
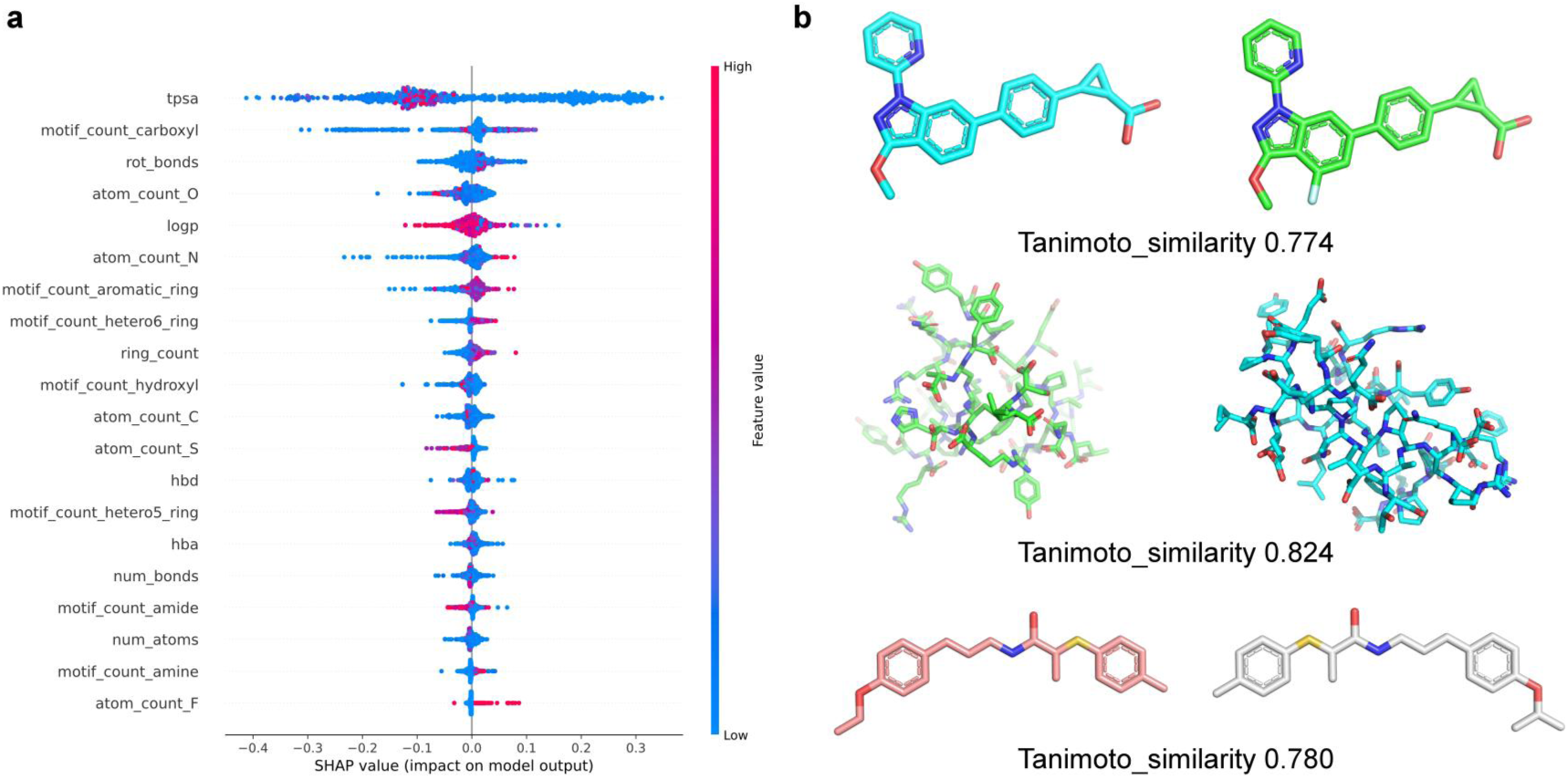
Multiscale interpretability via SHAP and counterfactual analysis. (a) Global feature attribution. SHAP identifies TPSA, lipophilicity, and hydrogen-bonding capacity (HBD/HBA) as primary determinants of BBB permeability. Atomic importance is color-coded from pink to blue, representing the contribution to positive prediction. (b) Visualization of physicochemical shifts for three representative pairs (Cases 1-3). Each trajectory illustrates the minimal structural and descriptor modifications required to flip the prediction from the seed state (BBB+, left) to the counterfactual state (BBB-, right).

The model consistently responds to chemically meaningful perturbations. In a small-molecule pair (Case 1), removing a single fluorine substituent from an aromatic ring flipped the prediction from BBB+ to BBB-. This shift is accompanied by a decrease in lipophilicity (Δ LogP = −0.14) despite nearly unchanged polarity, indicating that the model captures the contribution of halogenation to membrane partitioning.

In complex peptidomimetic structures (Case 2), prediction shifts are dominated by cumulative and context-dependent polarity effects. The model’s predictions diverge from conventional medicinal chemistry heuristics. In this peptidomimetic pair, the counterfactual (predicted BBB-) shows lower polarity (Δ TPSA = −56.0 Å²) and higher lipophilicity (ΔLogP = +1.01) than the seed molecule (predicted BBB+). While classical rules such as Lipinski^30^ and Veber^31^ would associate such shifts with enhanced permeability, our model predicts the opposite flip.

Structural inspection reveals reorganization of charged side chains and backbone amide environments, not just global descriptor shifts. The model captures local sub-structural patterns that serve as topological proxies for favorable 3D orientations. In peptide-like systems, permeability depends not only on overall polarity but also on how polar and charged groups are spatially arranged.

In Case 3, converting a linear ethoxy group (-OCH_2_CH_3_) to a branched isopropoxy substituent (-OCH(CH_3_)_2_) resulted in a discernible uplift in predicted BBB permeability from 0.48 to 0.55. Beyond alkyl branching, the counterfactual compound introduces a chiral center, where the model demonstrated acute sensitivity to stereochemical configurations.

These observations underscore that prediction shifts arise from context-specific structural modifications, consistent with global SHAP analysis where polarity and lipophilicity predominate.

### Scaffold-Level Misclassification Analysis

Evaluating how a predictive model responds to small structural changes in localized analog series is critical for its usefulness in lead optimization. By analyzing closely related congeners, we identified three scaffold families in which minor structural modifications cause substantial changes in experimental BBB permeability that the model does not consistently capture (**Supplementary Figure 5**).

One example is the aryl-sulfonyl series (**Supplementary Figure 5a**). While the model correctly classifies several non-permeable analogs, multiple experimentally validated BBB+ compounds containing para-difluoro, tert-butyl, or trifluoromethyl sulfonyl substituents are predicted as non-permeable, with positive probabilities below 0.05. Within these zones of high scaffold homogeneity, the model’s self-attention mechanism struggles to identify how terminal electronic withdrawing effects and localized steric bulk modulate the overall dipole moments and desolvation energies.

A related inconsistency occurs in the substituted benzamide series (**Supplementary Figure 5b**). Several mono- and di-chloro-substituted benzamides, experimentally classified as BBB+, are misclassified as BBB- with low predicted probabilities (pBBB+ 0.08-0.48). In contrast, closely related trifluoromethyl-substituted analogs are correctly predicted as permeable with much higher confidence (pBBB+ 0.93-0.96). Given the strong structural similarity across the series, these observations resemble activity-cliff-like behavior, where small substituent changes lead to large differences in predicted permeability.

Similar patterns are also observed in a side-chain-varied analog series (**Supplementary Figure 5c**), which comprises three closely related compounds differing primarily in peripheral aliphatic substituents. The model correctly predicts two experimentally validated BBB- compounds, but the remaining BBB+ analog is assigned similarly low predicted probability. This suggests a limitation of the 2D node-embedding framework, in which the influence of distal nonpolar extensions becomes progressively attenuated over successive message-passing layers. The resulting structural smoothing collapses latent variance, making embeddings insensitive to peripheral steric changes and ultimately preventing the model from distinguishing sharp permeability differences among closely related analogs.

### Mechanistic Case Studies and Model Limitations

To evaluate the practical utility of BBBP-Atlas and to examine the boundaries of structure-only modeling, we analyzed model predictions on a vetted set of 24 clinically characterized compounds (**Supplementary Table 5**) with well-established BBB permeability profiles, independent of the training data. This set includes FDA-approved CNS-active drugs, peripherally restricted therapeutics, as well as biologics and peptides with diverse and clearly defined BBB transport mechanisms.

For compounds whose BBB permeability is largely determined by intrinsic physicochemical properties, model predictions are consistent with clinical observations (**Supplementary Figure 6**). CNS-active drugs such as lorlatinib^32,33^ (BBB+, 96.8%), alectinib^34^ (BBB+, 90.7%), and levetiracetam^35^ (BBB+, 99.8%) are correctly classified with high confidence. They typically possess physicochemical profiles optimized for passive diffusion, specifically moderate molecular weight, low polar surface area, and limited hydrogen-bonding capacity within a more stringent range than general therapeutics.^36^ Conversely, large and highly polar antibiotics, including glycopeptides and aminoglycosides (e.g., vancomycin,^37^ BBB-, 98.4%; gentamicin,^38^ BBB-, 100%) are consistently predicted as impermeable under intact, non-inflamed BBB conditions, capturing the physicochemical cutoff beyond which passive diffusion becomes negligible. Thus, BBBP-Atlas effectively captures high-dimensional physicochemical determinants of passive permeability.

Discrepancies arise for compounds whose brain exposure depends on active transport processes (**Table 2**). The model misclassifies peptides that cross the BBB via receptor-mediated transcytosis (RMT), such as Angiopep-2^39^ (BBB-, discordant, 69.5%) and TfR-T12^40^ (BBB-, discordant, 98.7%). Although these peptides penetrate the brain in vivo through LRP1 or transferrin receptor engagement, they lack physicochemical features indicative of passive diffusion. As a result, the structure-only model underpredicts their permeability, highlighting a fundamental limitation in capturing receptor-mediated transport mechanisms.^41^ A similar pattern is observed for EGFR inhibitors such as osimertinib and gefitinib, whose brain exposure is modulated by context-dependent efflux.^42^ These cases indicate that permeability mediated by transporters, receptor engagement, or intracellular trafficking cannot be reliably inferred from molecular topology alone.

**Table 2.**
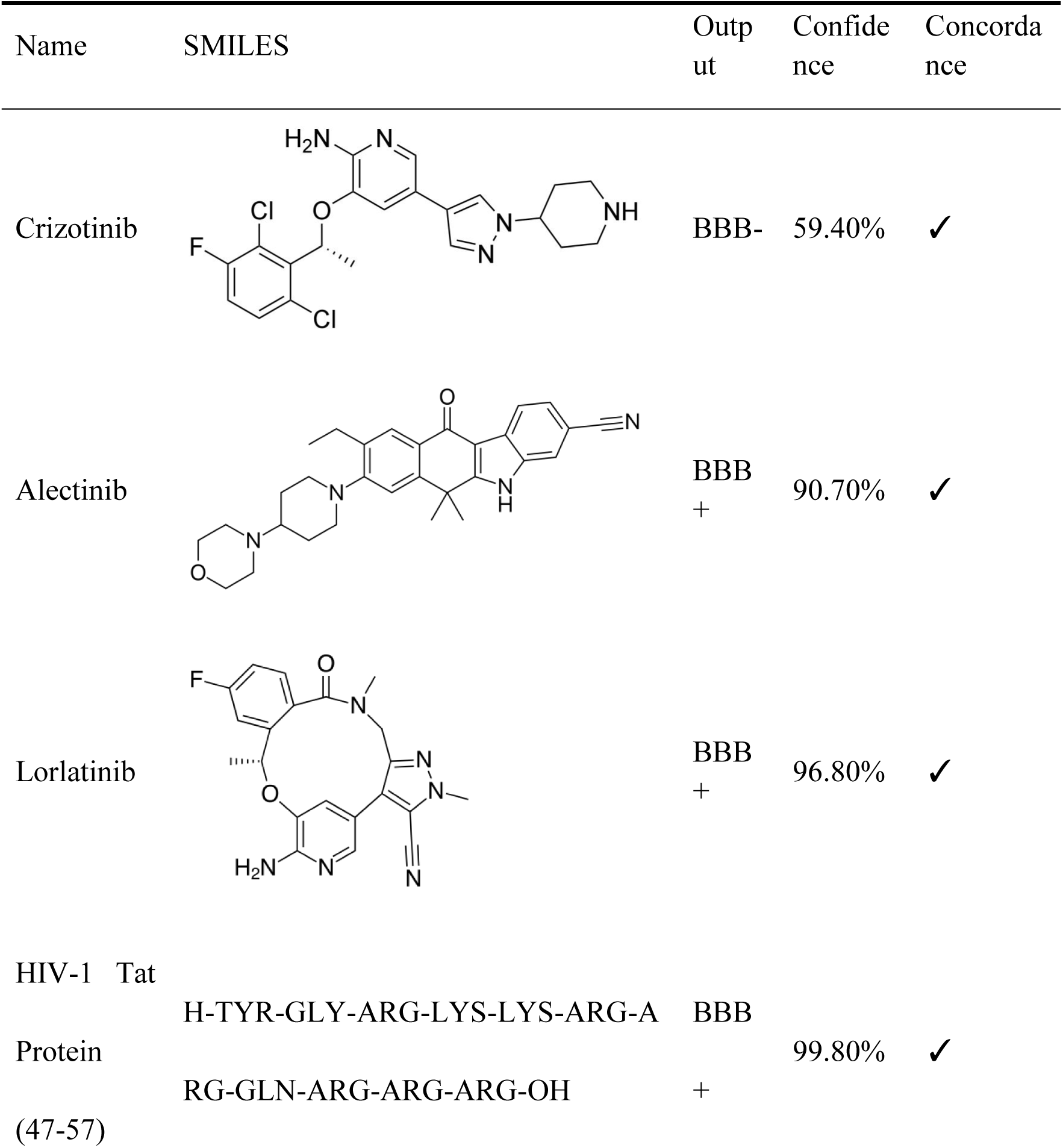
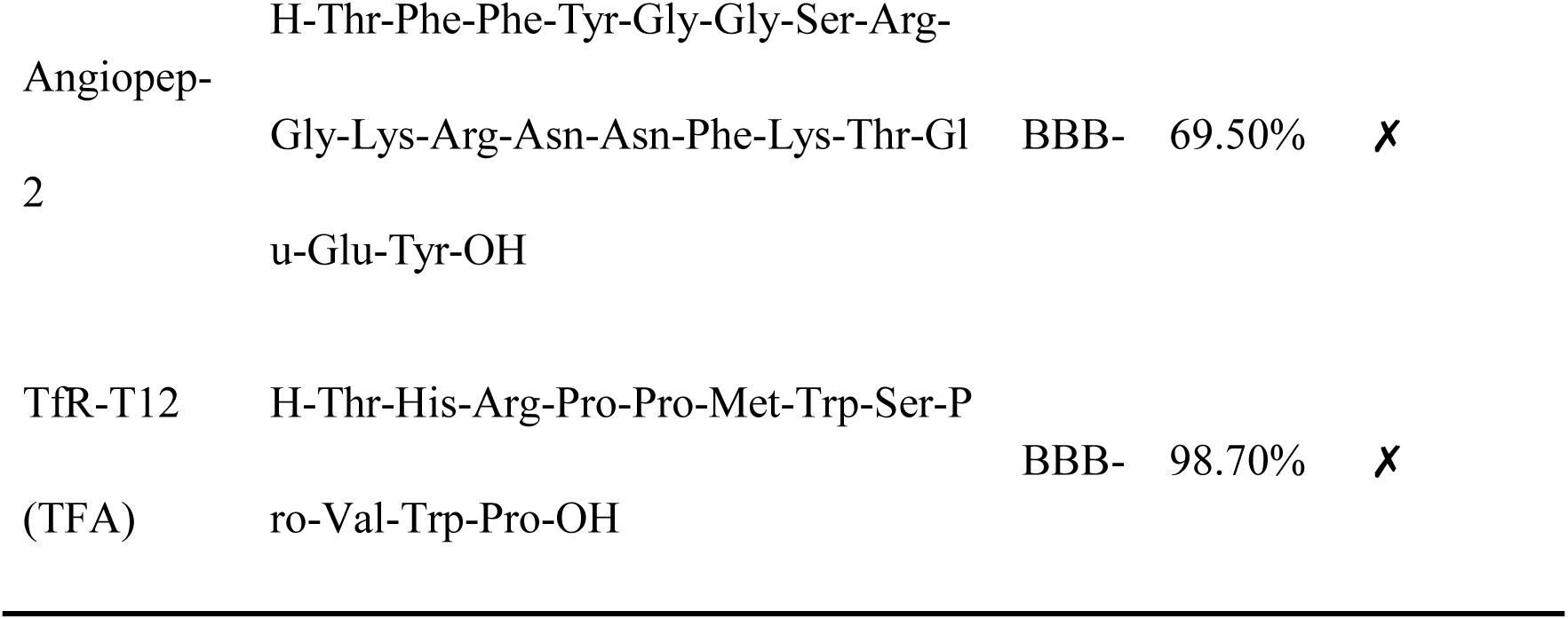
Representative case studies of model predictions on clinically characterized compounds. Clinically characterized compounds with known BBB permeability were evaluated using the proposed model. For each compound, the predicted class (BBB+ or BBB-), confidence score, and concordance with reported clinical permeability are shown. The full set of 24 compounds is provided in Supplementary Table 5.

| Name | SMILES | Output | Confidence | Concordance |
| --- | --- | --- | --- | --- |
| Crizotinib |  | BBB- | 59.40% | ✓ |
| Alectinib |  | BBB+ | 90.70% | ✓ |
| Lorlatinib |  | BBB+ | 96.80% | ✓ |
| HIV-1 Tat Protein (47-57) | H-TYR-GLY-ARG-LYS-LYS-ARG-ARG-GLN-ARG-ARG-ARG-OH | BBB+ | 99.80% | ✓ |
| Angiopep-2 | H-Thr-Phe-Phe-Tyr-Gly-Gly-Ser-Arg-Gly-Lys-Arg-Asn-Asn-Phe-Lys-Thr-Gl<br>u-Glu-Tyr-OH | BBB- | 69.50% | ✗ |
| TfR-T12<br>(TFA) | H-Thr-His-Arg-Pro-Pro-Met-Trp-Ser-P<br>ro-Val-Trp-Pro-OH | BBB- | 98.70% | ✗ |

BBBP-Atlas shows strong predictive performance when BBB permeability is governed by intrinsic molecular structure, while its reduced reliability for RMT-dependent peptides and efflux-substrate drugs defines the practical scope of structure-based modeling. High-confidence predictions are therefore most reliable for passive diffusion, whereas transporter- or receptor-dependent compounds should be interpreted with caution and warrant additional experimental validation.

### Online platform

An interactive web platform for BBBP-Atlas is publicly available at https://cadd.drugflow.com/bbbp/, providing open-access BBB permeability prediction. The platform is deployed using a Django-based backend architecture, with curated BBB data stored in an optimized SQLite relational database to support efficient querying and scalable online inference. The frontend is engineered via HTML, CSS, and JavaScript, providing a responsive interface for compound browsing, dataset exploration, and online prediction, as shown in **Figure 6**.

**Figure 6.**
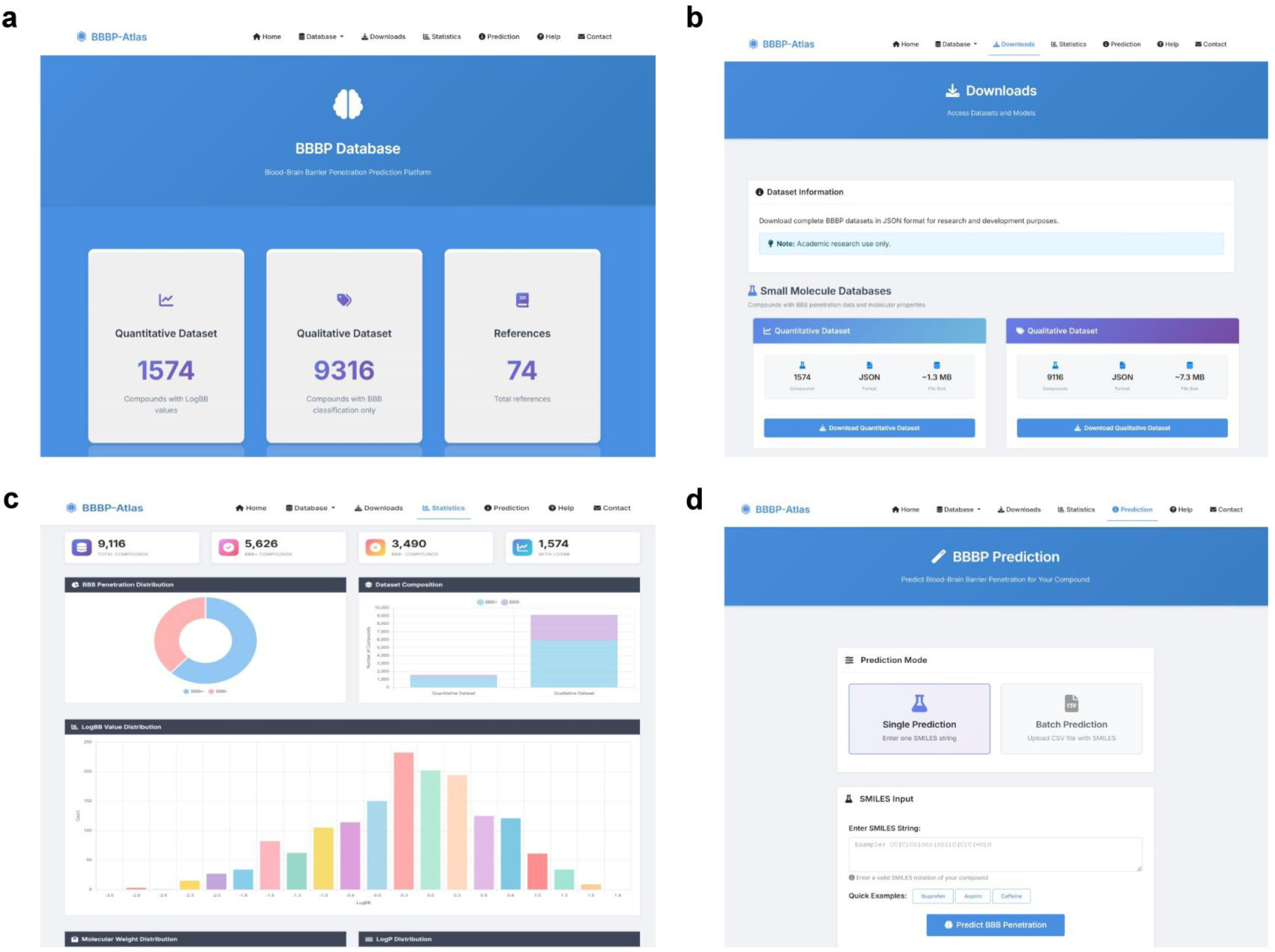
Architecture and functional modules of the BBBP-Atlas web platform. (a) Unified navigation framework enabling access to Home, Database, Downloads, Statistics, Prediction, Help, and Contact. (b) Data repository overview, including 9,316 small molecules and 902 peptides, and a quantitative subset of 1,574 compounds with logBB values. (c) Statistics module with interactive visualization of chemical space and property distributions. (d) Prediction interface supporting single SMILES input and batch CSV upload, with outputs including BBB classification and confidence scores.

The web server integrates the pretrained BBBP-Atlas model, which achieves 85% accuracy on external benchmark datasets. Users can submit molecules via two modes. In the single-compound submission track, users input molecular structure as a SMILES string. The backend dynamically parses the textual syntax into a 2D topological molecular graph, executing forward-propagation inference to output binary permeability assessments (BBB+/BBB-). In the batch prediction mode, users may upload CSV files containing a designated “SMILES” column. The backend parallelizes the processing of molecular libraries, optimizing computational throughput and resource allocation during large-scale virtual screening workflows. All predictions are returned with a binary label and a confidence score, with results downloadable as CSV for further chemoinformatics analysis. User documentation and example workflows are also provided to facilitate platform accessibility and reproducibility.

## DISCUSSION

In this study, we proposed OmniBBBP, a non-redundant dataset comprising 10,218 compounds (9,316 small molecules and 902 peptides), and developed a cross-modal framework that achieves strong predictive performance on independent test sets (ACC = 0.8914, MCC = 0.7678), with robust generalization to an external balanced dataset (AUC = 0.9108, ACC = 0.85). While prior BBBP predictive studies have been predominantly constrained to small molecules, BBBP-Atlas offers a broader scope by incorporating peptides into a unified modeling framework. By exploiting cross-modal knowledge transfer, the model effectively mitigates the data disparity between the two subsets, allowing the structural patterns learned from large-scale small molecule data to enhance peptide permeability predictions.

Our analysis reveals a fundamental limitation of structure-only modeling. The model reliably captures intrinsic BBB permeability governed by physicochemical properties, as illustrated by its correct discrimination between crizotinib (BBB-, 59.4%) and alectinib and lorlatinib (BBB+, >90%). However, its performance deteriorates for compounds whose brain exposure depends on active transport processes, including efflux-regulated drugs (e.g., osimertinib, gefitinib) and receptor-mediated peptides (e.g., angiopep-2). While passive diffusion is readily learnable from molecular topology, our results indicate that structure-based representations alone may be insufficient for capturing active transport mechanisms without additional biological contextualization or targeted, mechanism-specific training data.

This limitation suggests that purely structure-based architectures may approach a performance ceiling. Instead, future progress will depend on integrating molecular representations with mechanistically relevant biological context, including explicit annotations of transporter substrates (e.g., P-gp, BCRP) and receptor-mediated pathways. BBBP-Atlas provides both a high-quality benchmark and a mechanistic framework to disentangle structure-driven and biologically regulated permeability, supporting the rational design of CNS-active therapeutics and agents targeting brain metastases.

## METHODS

### Dataset Construction

We developed OmniBBBP, a curated collection of experimentally validated BBBP annotations for small molecules and peptides. Data were retrieved from PubMed and Google Scholar using the keywords (“blood-brain barrier permeability” or “BBB permeability” or “logBB”) and (“small molecule” or “peptide”), covering studies from January 2000 to December 2025. This search was supplemented with domain-specific databases, including B3Pdb^22^ and Brainpeps.^43^

To ensure data quality, a two-step curation protocol was applied. First, only entries with explicit molecular representations (SMILES for small molecules, FASTA for peptides) and unambiguous permeability annotations were retained. For categorical labels, we standardized all positive annotations (e.g., “BBB+”/“permeable”/“CNS+”) as 1 and negative annotations as 0. Quantitative logBB values were binarized using a threshold of −1 (logBB > −1 for BBB+, otherwise BBB-), following commonly adopted thresholds in prior BBBP studies.^44,45^ Second, for compounds with conflicting annotations across sources, we prioritized peer-reviewed experimental measurements over computational predictions and, if conflicts persisted, excluded the entry. All retained structures were standardized using RDKit^46^ (v2025.9.2), including salt removal, solvent stripping, and hydrogen normalization, while preserving stereochemistry.

The final OmniBBBP comprises 10,218 unique entries, including 9,316 small molecules and 902 peptides (**Supplementary Figure 1**) from 74 publicly available sources. Among these, 1,574 compounds have experimentally reported logBB values; these were used for quantitative benchmarking and binarized for consistency with the primary classification task. The peptide collection comprises 732 standard peptides and 170 sequences containing non-natural or chemically modified residues. Among BBB+ peptides, 22.1% contain modified residues, compared with 15.6% in the BBB-subset. Analysis of the clean peptide sequences further showed that BBB+ peptides were generally shorter, with an average length of 15.61 residues versus 17.04 residues for BBB- peptides, and also exhibited a substantially higher average net charge, reaching +2.48 compared with +0.84 for BBB- peptides. In contrast, Chou-Fasman secondary-structure propensity scores were comparable between the two groups, with average values close to 1.0.

Compared to the widely used MoleculeNet BBBP dataset^21^ (2,053 compounds, primarily small molecules), OmniBBBP is approximately five times larger and uniquely incorporates 902 peptides, addressing a critical gap in peptide-based BBB permeability modeling.

### Backbone

We designed and developed the BBBP-Atlas backbone tailored to decode complex molecular topologies and peptide cross-modal interactions. While graph-based attention networks have shown therapeutic promise in general structural prediction tasks,^47^ our architecture explicitly reformulates these attention principles to optimize dynamic structural learning specifically for blood-brain barrier penetration modeling. By incorporating an advanced self-attention mechanism, our model captures multi-scale, long-range relationships between non-adjacent nodes, enabling highly contextualized weighting of neighboring atoms during message passing. In the input molecular graph G_p_ = (H_p_, E_p_, X_p_), the node features 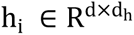 for the i-th node and edge features 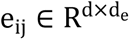 for the edge connecting node i and node j are initialized to 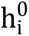 and 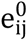, respectively, in a d-dimensional space using two linear layers.

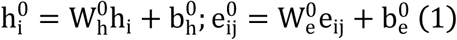

where 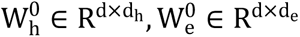, and 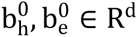. After passing through the embedding layer, the features are expanded to the same dimension. Then, message passing and aggregation are executed via the convolutional layers. In the BBBP-Atlas, this operation is primarily achieved by the self-attention mechanism. The model performs six convolutional operations, with the convolution process for the l-th layer expressed by the following equations:

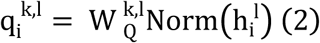

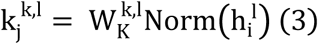

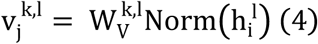

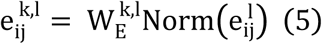

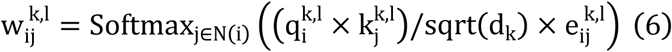

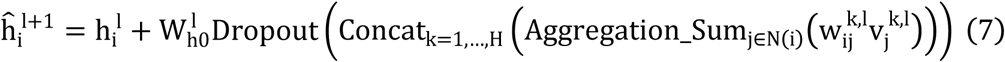

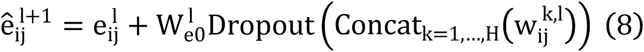

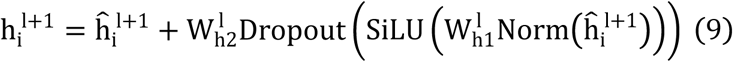

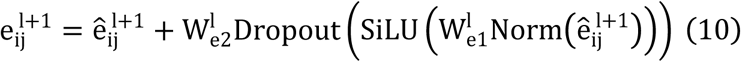

where 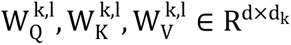 and 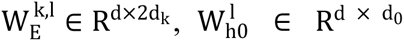 and 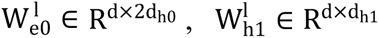 and 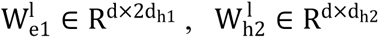 and 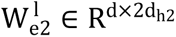 represent learnable parameters derived from linear layers. k ∈ 1, …, H signifies the number of attention heads, while d_k_ represents the dimension of each head, calculated as d divided by H. j ∈ N(i) denotes the neighboring nodes of node i, with Norm indicating batch normalization. Concatenation is denoted by Concat, and Dropout denotes the dropout operation. Activation functions are represented by SiLU.

The aggregation of messages on the edges connecting node i and its neighboring nodes j is depicted by Aggregation_Sum_j∈N(i)_, while Softmax_j∈N(i)_ symbolizes the SoftMax operation applied to the neighboring nodes j.

### Model training

The 4-layer, 4-head BBBP-Atlas model described above was trained on an 8:1:1 train/validation/test split. Molecular graphs were constructed with nodes representing atoms (38-dimensional features encoding atom type, degree, formal charge, radical electrons, hybridization, aromaticity, and hydrogen counts) and edges representing bonds (6-dimensional features capturing bond type, conjugation, and ring membership). Feature normalization followed the procedure described in the Backbone section. SMILES strings for small molecules were standardized with salts and solvents removed, while peptide FASTA sequences were length-adjusted and encoded consistently.

Training was performed using the Adam optimizer^48^ with a learning rate of 5×10^−4^ and batch size of 32 for up to 300 epochs. Binary cross-entropy loss with logits was used, with SiLU activation and a dropout rate of 0.1 applied after each attention layer. A ReduceLROnPlateau scheduler (factor 0.9, patience 3) adjusted the learning rate based on validation loss, and early stopping was triggered if validation loss did not improve for 10 consecutive epochs. Batch normalization was applied to stabilize training.^49^

Model checkpoints were saved at each epoch, and the model achieving the highest validation MCC was selected for final evaluation. All experiments were conducted on NVIDIA H800 GPUs, with fixed random seed set to 88 to ensure reproducibility. Training on the full dataset required approximately 1 hour per fold.

### Benchmark Metrics

Model performance in BBB permeability prediction was evaluated using seven complementary metrics: area under the ROC curve (AUC), accuracy (ACC), F1-score (F1), Matthews correlation coefficient (MCC), balanced accuracy (BA), sensitivity (SE), and specificity (SP). These metrics were selected to address three specific challenges in the domain: (i) class imbalance, MCC and BA are robust to skewed class distributions, unlike raw ACC; (ii) asymmetric error costs, in CNS drug screening, false negatives (missing a permeable compound) are more costly than false positives, making the SE/SP trade-off essential to report; and (iii) threshold dependence, AUC provides a threshold-independent measure of ranking quality.

Accordingly, MCC serves as our primary metric due to its dependence on all four confusion matrix categories^50^ (TP, TN, FP, FN) and its insensitivity to class imbalance. SE and SP characterize the clinically relevant trade-off, while F1 balances precision and recall. BA and ACC provide overall correctness checks, and AUC quantifies the probability that a randomly selected BBB+ compound ranks higher than a randomly selected BBB- compound. Mathematical definitions are provided below.

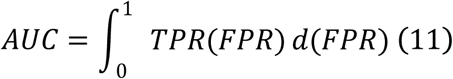

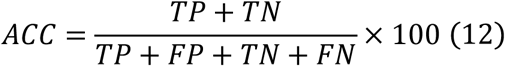

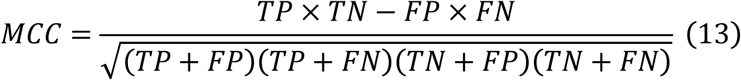

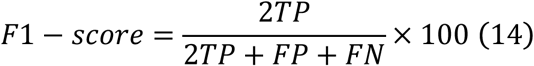

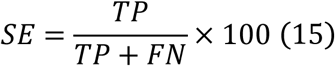

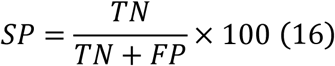

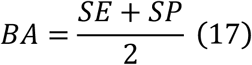

## ASSOCIATED CONTENT

The datasets utilized in our study were manually curated and standardized from publicly available sources. The complete processed dataset and associated materials have been deposited in Zenodo (https://doi.org/10.5281/zenodo.20158745). The code used in the study is publicly available from the GitHub repository (https://github.com/SHENXIN516/BBBP-Atlas).

## ACKNOWLEDGMENT

This work was supported by grants from the “Pioneer” and “Leading Goose” R&D Program of Zhejiang (2025C01117), the National Key R&D Program of China (2024YFA1307501), and Central Guidance for Local Science and Technology Development Funds Project (2025ZY01022).

## ABBREVIATIONS

BBB: Blood-brain barrier
BBBP: Blood-brain barrier permeability

## Notes

https://doi.org/10.5281/zenodo.20158745

### Competing Interest Statement

The authors have declared no competing interest.

